# Chimpanzee SIV Envelope trimer: structure and deployment as an HIV vaccine template

**DOI:** 10.1101/459164

**Authors:** Raiees Andrabi, Jesper Pallesen, Joel Allen, Ge Song, Jinsong Zhang, Natalia de Val, Gavin Gegg, Katelyn Porter, Ching-Yao Su, Matthias Pauthner, Amanda Newman, Hilary Bouton-Vervelle, Fernando Garces, Ian A. Wilson, Max Crispin, Beatrice H. Hahn, Barton F. Haynes, Laurent Verkoczy, Andrew B. Ward, Dennis R. Burton

## Abstract

Epitope-targeted HIV vaccine design seeks to focus antibody responses to broadly neutralizing antibody (bnAb) sites by sequential immunization. Chimpanzee SIV Envelope (Env) shares a single bnAb site, the V2-apex, with HIV, suggesting its possible utility in an HIV immunization strategy. Accordingly, we generated a chimpanzee SIV Env trimer, MT145K, which displays selective binding to HIV V2-apex bnAbs and precursor versions, but no binding to other HIV specificities. We determined the structure of the MT145K trimer by cryo-EM and showed its architecture was remarkably similar to HIV Env. Immunization of an HIV V2-apex bnAb precursor Ab-expressing knock-in mouse with chimpanzee MT145K trimer induced HIV V2-specific neutralizing responses. Subsequent boosting with an HIV trimer cocktail induced responses exhibiting some virus cross-neutralization. Overall, the chimpanzee MT145K trimer behaves as expected from design both in vitro and in vivo and is an attractive potential component of a sequential immunization regimen to induce V2-apex bnAbs.

## Introduction

The ability to induce human immunodeficiency virus (HIV) envelope (Env) specific broadly neutralizing antibodies (bnAbs) will likely be a key feature of a prophylactic vaccine immunogen. Potent Env-specific bnAbs are produced in a small subset of HIV infected donors, yet attempts to elicit such responses through immunization have failed to date (Escolano et al., 2017; Haynes and Mascola, 2017; McCoy and Burton, 2017; Ward and Wilson, 2017). Previous studies have revealed that the HIV bnAb germline reverted precursors possess unique features that greatly reduce their overall frequencies in the B cell immune repertoire and, hence, their ability to be targeted by vaccines (Briney et al., 2012; Haynes et al., 2012; Kepler et al., 2014; Klein et al., 2013; Verkoczy et al., 2010; Xiao et al., 2009). Therefore, recent immunogen design approaches that seek to induce bnAb responses by vaccination are taking these rare precursor features into consideration to efficiently activate bnAb precursors and shepherd them along favorable bnAb developmental pathways (Andrabi et al., 2015; Escolano et al., 2016; Gorman et al., 2016; Jardine et al., 2013; McGuire et al., 2013; Saunders et al., 2017; Steichen et al., 2016a). This design approach has shown great promise for two of the HIV Env bnAb sites, namely the CD4 binding site (CD4bs) and the V3-N332 glycan site in animal models expressing the appropriate germline precursors (Andrabi et al., 2018; Briney et al., 2016; Dosenovic et al., 2015; Escolano et al., 2016; Jardine et al., 2015; McGuire et al., 2016; Sok et al., 2016a; Steichen et al., 2016b; Tian et al., 2016; Williams et al., 2017). Thus, immunogen designs and strategies that can select rare bnAb precursors and reduce off-target B cell responses are valuable for nAb immunofocusing efforts.

One of the Env sites that has shown great promise for vaccine targeting is the V2 apex bnAb epitope (Andrabi et al., 2015; Gorman et al., 2016; Voss et al., 2017). This bnAb epitope sits at the 3-fold axis of the trimer and is primarily formed by a patch rich in positively charged lysine residues and protected by two glycans at HXB2 HIV-1 reference positions N160 and N156/N173 that are part of the Env glycan shield (Andrabi et al., 2017; Bhiman et al., 2015; Bonsignori et al., 2011; Doria-Rose et al., 2014; Gorman et al., 2016; Julien et al., 2013b; Lee et al., 2017; McLellan et al., 2011; Pancera et al., 2013; Walker et al., 2011; Walker et al., 2009). The bnAb precursors targeting this site possess a long anionic heavy-chain complementarity-determining region 3 (CDRH3) that penetrates the glycan shield to reach the protein epitope surface underneath (Bonsignori et al., 2011; Doria-Rose et al., 2014; Landais et al., 2017; Lee et al., 2017; McLellan et al., 2011; Walker et al., 2011; Walker et al., 2009). BnAb prototypes within this class interact with the V2 apex bnAb protein-glycan core epitope through common germline-encoded motifs and are, thus, targetable by unique trimers that bind with their germline Ab versions, previously reported by us and others (Andrabi et al., 2015; Gorman et al., 2016). Hence, the germline-priming immunogens to this site could be based directly on native-like trimer configurations (Sanders et al., 2013; Sanders et al., 2015). Other features that favor this site for vaccine targeting are: a) V2 apex bnAbs are elicited frequently in humans that make bnAbs, b) they emerge early in infection, and c) they possess relatively low levels of somatic mutation compared to most other HIV Env bnAbs (Bonsignori et al., 2011; Doria-Rose et al., 2014; Georgiev et al., 2013; Kepler et al., 2014; Landais et al., 2016; Landais et al., 2017; Moore et al., 2011; Walker et al., 2009; Wibmer et al., 2013).

Of note, among the major HIV Env bnAb specificities that include V2-apex, V3-N332, CD4bs and gp120-41 interface, the V2 apex site is the only bnAb site that consistently exhibits cross-group neutralizing activity with virus Envs derived from HIV-1 group M, N, O and P (Braibant et al., 2013; Morgand et al., 2016). In addition, V2-apex bnAbs display cross-neutralizing activity with the Simian Immunodeficiency Virus (SIV) isolates that infect chimpanzees (SIVcpz*Ptt* [*Pan troglodytes troglodytes*], SIVcpz*Pts* [*Pan troglodytes schweinfurthii*]) and gorillas (SIVgor) (Barbian et al., 2015). The retention of the V2 apex bnAb epitope at the time of species cross-over from chimpanzees to humans highlights the biological significance of this region and here we sought to design a trimer based on the SIVcpz*Ptt* Env sequence that could potentially guide an immunofocused response to the HIV V2 apex bnAb epitope. We hypothesized that an SIVcpz*Ptt*-based trimer will not only help to specifically enrich V2 apex-specific B cells but also, owing to V2 apex species cross-conservation, could help guide a V2-focused nAb response when coupled with HIV trimers in a sequential prime/boost immunization strategy. In such an immunization scheme, overall Env backbone sequence diversity in combination with conservation of the V2 apex bnAb epitope in sequentially administered immunogens is likely to reduce germinal center competition for V2 apex-specific B cells (Tas et al., 2016; Wang et al., 2015). Such a scheme could not only favor a B cell recall response to the V2 apex bnAb epitope but also reduce off-target Env-specific responses.

We designed here an SIVcpz*Ptt*-based trimer, MT145K, that displays native trimer-like properties, and selectively binds V2 apex bnAbs as well as their germline reverted precursor versions. We determined a structure of the MT145K trimer by cryo-EM at a global resolution of 4.1Å and the overall architecture was remarkably similar to HIV Env trimers (Julien et al., 2013a; Lyumkis et al., 2013; Ozorowski et al., 2017; Pancera et al., 2014). In addition, the glycan shield composition of MT145K closely resembled that of HIV Env glycans but was sufficiently different in positioning of the glycans to exclude binding of all HIV bnAbs except for those directed to the V2 apex. MT145K trimer immunization in a V2 apex unmutated common ancestor (UCA)-expressing knock-in mouse model revealed induction of a predominantly V2-apex-site neutralizing Ab response that was reproducible and cross-neutralized a related set of HIV isolates. Overall, the chimpanzee MT145K immunogen shows promise as an immunogen in HIV vaccination strategies.

## Results

### Selection and design of a chimpanzee Env-derived trimer

Immunogen templates based on native-like Env trimers offer great potential for HIV vaccine development, as they display bnAb epitopes and largely occlude non-native epitopes. However, it remains challenging to induce an epitope-focused bnAb response with Env trimer immunogens, as the bnAb epitopes are relatively immunoquiescent and even very limited exposure of non-desirable epitopes can disturb responses to bnAb epitopes (Havenar-Daughton et al., 2017; Wang et al., 2015). Therefore, trimer designs and/or strategies that can mask non-relevant immunodominant epitopes or reduce induction of off-target Ab responses could help guide immunofocused neutralizing responses. In addition, typical lack of interaction of Env forms with germline-reverted bnAb precursors means difficulties in activating the appropriate B cell lineages. Accordingly, we undertook design of a trimer immunogen that could help guide an epitope-focused Ab response to the V2 apex site of HIV Env. Based on previous studies, we hypothesized that a chimpanzee SIVcpz*Ptt/Pts* or gorilla SIVgor Env sequence-based trimer that shares the V2 apex bnAb epitope with HIV-1 could enrich B cell precursors and boost responses specific to this site (Barbian et al., 2015). Since SIVcpz*Ptt*, among various SIV-species Env sequences, are phylogenetically closest to the HIV-1 Env, we surmised that the SOSIP.664 trimer-stabilizing modifications, which have been used on several HIV-1 Env backgrounds, could be employed for stabilization of soluble SIV Env (Gao et al., 1999; Sanders et al., 2013; Sharp and Hahn, 2011).

We incorporated the SOSIP.664 trimer design modifications into four SIVcpz*Ptt* Env sequences; GAB1, MB897, EK505, and MT145 (Figure S1A). These isolates have been previously shown to be sensitive to the V2 apex bnAbs, PG9, PG16, and PGT145 (Barbian et al., 2015). Further characterization showed that one of these SIVcpz*Ptt* Env sequences, MT145 SOSIP.664, could be expressed as a soluble Env trimer protein (Figure S1B). PGT145 Ab affinity-purified MT145 trimer was efficiently cleaved into gp120 and gp41 subunits, and revealed well-ordered native-like trimer configurations that were highly thermostable, which are all properties displayed by natively folded HIV-1 soluble trimers (Figure S2A-D) (Pugach et al., 2015; Sanders et al., 2013; Sharma et al., 2015).

### MT145K trimer binds prototype V2 apex bnAb precursors

One property thought to be critical for vaccine immunogens to select rare bnAb precursors is the ability to effectively bind to UCA B cell receptors (Dosenovic et al., 2015; Escolano et al., 2016; Jardine et al., 2015; McGuire et al., 2016; Steichen et al., 2016a). Therefore, to gain or improve binding of the V2 apex bnAb inferred precursor Abs to MT145 Env trimer, we substituted a glutamine (Q) with a lysine (K) residue (HXB2 position 171) in strand C of the V2 apex bnAb core epitope (Figure 1A-B). We based this substitution on the presence of a positively charged motif (KKKK) in CRF250 and CP256.SU strand C V2 Env sequences, both of which bind V2 apex bnAb prototype precursors (Andrabi et al., 2015; Doria-Rose et al., 2014; Gorman et al., 2016). ELISA binding revealed strong binding of the mature V2 apex bnAb prototypes with the MT145-WT trimer and weak but detectable binding with one of the UCA Abs, CAP256 UCA (Figure 1C). Strikingly, binding with our V2-engineered MT145 trimer (henceforth referred to as “MT145K”) not only improved binding to CAP256 UCA Ab but also conferred binding on both PG9 and CH01 iGL Abs (Figure 1C). The PG9 and CH01 iGL Abs used here had diversity (D; heavy chain) and joining (J; both heavy and light chains) genes reverted to their corresponding germline gene families in the CDRH3s, in addition to the VH and VL regions reported previously (Figure S3) (Andrabi et al., 2015; Gorman et al., 2016).

**Figure 1.**
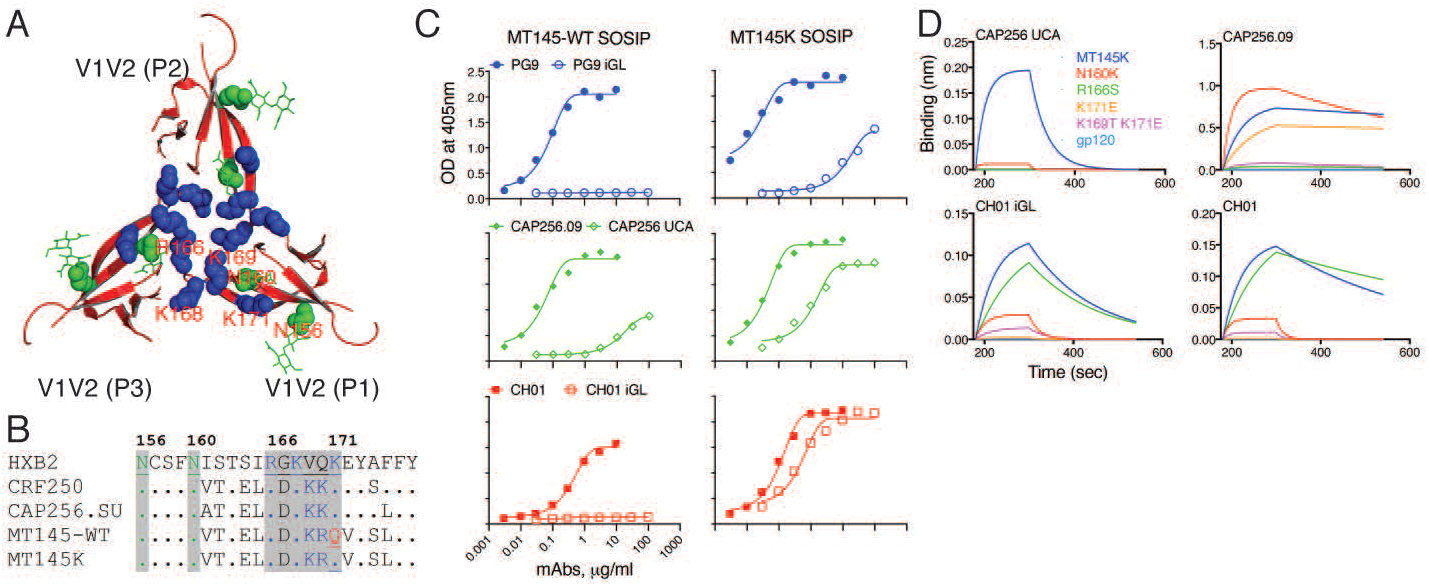
Design of a chimpanzee Env-stabilized trimer and binding to V2 apex bnAb iGL Abs. **A.** Structural arrangement of the V2 apex bnAb core epitope region on BG505.664 soluble Env trimer (modified from (Garces et al., 2015) (PDB: 5CEZ)). The ribbon representation of V1V2 loop strands that form the trimer apex show a cluster of positively charged lysine-rich peptide regions (HXB2-R166-K171: R or K residues shown as blue spheres) and the two glycans N156 and N160 (depicted in green spheres with lines). The side chains of the positively charged residues intersperse with the side chains of residues from adjacent protomers to form a continuous positively charged surface at the tip of the trimer to provide a minimal V2 apex bnAb epitope. **B.** Amino-acid sequence alignment of strand B and C V2 of HIV CRF250, CAP256.SU, chimpanzee SIV MT145 WT and its V2-modified variant (Q171K), MT145K. Glutamine (Q) at position 171 (shown in red) was substituted with lysine (K) in MT145 Env to gain binding to V2 apex bnAb inferred germline (iGL) Abs. **C.** ELISA binding of mature V2 apex bnAbs, PG9, CAP256.09 and CH01 and their iGL versions to WT MT145 (red) and MT145K SOSIP trimers. **D.** Octet binding curves (association: 120s (180–300) and dissociation: 240s (300–540)) of CAP256 UCA and CH01 iGL Abs and their respective mature Ab versions (CAP256.09 and CH01) to MT145K trimer, its glycan knock-out (N160K) variant, K-rich core epitope substituted variants and the corresponding monomeric gp120. The Abs were immobilized on human IgG Fc capture biosensors and 1uM trimer or gp120 proteins used as analytes. The binding response is shown as nanometer (nm).

Previous mapping studies have defined the HIV core epitope recognized by mature V2 apex bnAbs (Andrabi et al., 2015; Gorman et al., 2016; Landais et al., 2017; Lee et al., 2017; McLellan et al., 2011; Pancera et al., 2013; Walker et al., 2009). To examine the contributions of V2 apex core epitope glycan and protein residues to binding by V2 apex bnAb inferred germline-reverted (iGL) Ab versions, we generated MT145K strand C peptide and glycan trimer variants that are known to eliminate interactions of V2 apex bnAbs with the Env trimer (Andrabi et al., 2015; McLellan et al., 2011; Pancera et al., 2013). Bio-Layer Interferometry (BLI or octet) binding analyses of the iGL Abs with these trimer variants showed that glycan/peptide epitope requirements of precursor Abs were largely similar to the requirements of corresponding mature Abs (Figure 1D), suggesting that most contacts with the MT145K V2 apex core epitope are already encoded in the germline configuration for this class of bnAbs. Notably, the mature Abs showed slightly more tolerance to changes within the core protein epitope, particularly for the CAP256.09 bnAb, suggesting that part of the affinity maturation in this class of Abs may be to accommodate variation within the bnAb V2 apex core epitope. Overall, the strand C V2-modification in the MT145 SOSIP.664 trimer conferred binding to multiple V2 apex bnAb germline prototypes.

### Architecture of the MT145K trimer

We solved the structure of the MT145K trimer by cryo-EM to a global resolution of ~4.1 Å (Figure S4, Table S1). Our structure represents the first atomic level structure of an SIV Env trimer. Like other class I fusion proteins, protomers (gp120 and gp41) of MT145K trimerize to form a metastable pre-fusion Env trimer (Figures 2A-B, S5). The trimer architecture exhibits a mushroom-like shape with subunits gp120 and gp41 constituting the envelope-distal and proximal entities, respectively (Figure 2B). Overall, the MT145K trimer configuration closely resembles that of the trimeric HIV-1 Env spike, with an overall Ca root mean square deviation (rmsd) of 1.9 Å (Kwon et al., 2015). Arrangement of V-loops in the MT145K Env trimer is reminiscent of the V-loop arrangement in the HIV Env trimer and is suggestive of a similar role in immune evasion by steric occlusion of underlying conserved epitopes (Julien et al., 2013a; Pancera et al., 2014). Notably, the V1 and V2 loops are largely solvent-exposed and occlude access to the underlying V3 loop (Figure 2C). Inaccessibility of the V3 loop is mediated by intra-protomer interactions of V1V2 to V3 and by extensive inter-protomer V1V2 trimer interactions at the apex of the spike. The SIV Env trimer exhibits well-ordered V2-V5 loops, while V1 is somewhat disordered (Figure 2D).

**Figure 2.**
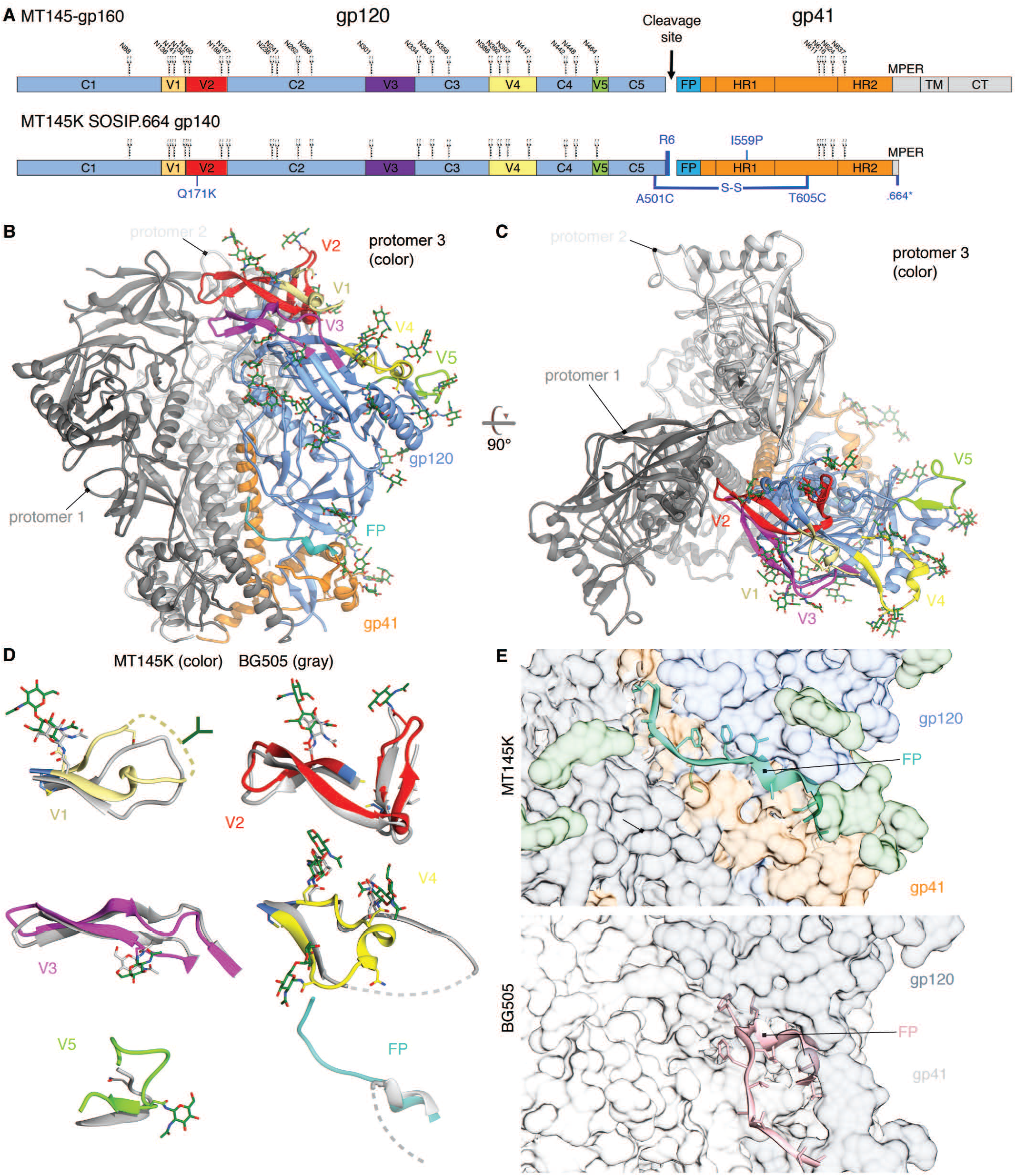
Cryo-EM structure of the MT145K trimer. **A.** Schematic showing MT145K SOSIP soluble trimer design from its full-length gp160 Env sequence. The gp120 constant (C1-C5) and variable (V1-V5) regions and the gp41 regions (fusion peptide (FP), heptad repeat (HR1 and HR2), membrane proximal external region (MPER), transmembrane (TM) and cytoplasmic tail (CT)) are indicated. The N-linked glycan positions for each NXT/S residue are labeled according to the HIV HXB2 numbering scheme. The SOSIP trimer stabilizing modifications include: (i) disulfide bond: A501C-T605C, (ii) R6 cleavage site, (iii) I559P, and (iv) 664-residue truncation in gp41 MPER. The substitution to incorporate a K-residue at position 171 (Q171K) to gain binding for V2 apex iGL Abs is indicated in blue. **B-C.** Side and top views of the unliganded MT145K trimer model based on the cryo-EM density map at ~4.1 Å resolution. Ribbon representations of the MT145K trimer spike, in which the subunits gp120 (cornflower blue) and gp41 (orange) are depicted on one protomer. The gp120 variable loops (V1-V5) positioned to the trimer periphery are depicted in different colors (V1: khaki, V2: red, V3: magenta, V4: yellow and V5: chartreuse). The fusion peptide region of gp41 is shown in cyan. Glycan sugar residues modeled based on density are represented in forest green stick form. **D.** Superimposition of variable loops (V1-V5) and fusion peptide region for MT145K and unliganded HIV clade A BG505 (PDB: 4ZMJ) SOSIP trimers. The dotted lines indicate regions in the V-loops or FP for which the observed electron density was absent or unclear. **E.** Structural comparison of gp41 regions of MT145K (orange) and BG505 (grey) trimers. The gp41 structural elements overall show a similar arrangement except for the fusion peptide region (colored cyan on MT145K and pink on BG505), that is exposed on the BG505 trimer but remains hidden in a pocket inside the MT145K trimer.

Proximal to the viral membrane is the gp41 subunit that forms the base of the trimer spike and is arranged into heptad repeat-1 (HR1), HR2 and the fusion peptide (FP) (Figure 2B). Similar to the HIV Env trimer, the three C-terminal helices of HR1 are centrally positioned along the trimer axis perpendicular to the viral membrane (Julien et al., 2013a; Lyumkis et al., 2013; Pancera et al., 2014). Intriguingly, the FP region, which has been observed solvent-exposed on the outside of the HIV-1 Env, is positioned in a pocket inside the MT145K trimer and remains sequestered in all three protomers (Figure 2E).

### Conservation of the glycan shield on HIV and SIV Env trimers

To compare the nature of the glycan shield on SIVcpz*Ptt* Env and HIV Env, we performed site-specific glycan analysis of the MT145K trimer. The overall oligomannose content of the MT145K trimer is similar to HIV Env (Figure 3A-B) (Panico et al., 2016; Pritchard et al., 2015) . However, although the distributions differed from the HIV-1 clade A strain BG505, which is dominated by Man_9_GlcNAc_2_ oligomannose-type glycans, MT145K is predominantly Man_8_GlcNAc_2_ (Behrens et al., 2016). In addition, further processing was evident in the MT145K trimer which showed elevated Man_6-7_GlcNAc_2_ structures (Figure 3B, S6). The outer domain of gp120 presents a high density of oligomannose glycans that form the intrinsic mannose patch (Bonomelli et al., 2011), which was a highly conserved feature across the two viral species. The apex of the MT145K trimer possessed oligomannose-type glycans at N160 that correspond to the trimer associated mannose patch (TAMP) also observed on HIV-1 Env (Behrens et al., 2017). As for HIV-1, glycans at the base of the trimer at N88 and on gp41 of the MT145K trimer were extensively processed (Figures 3A, C, S6).

**Figure 3.**
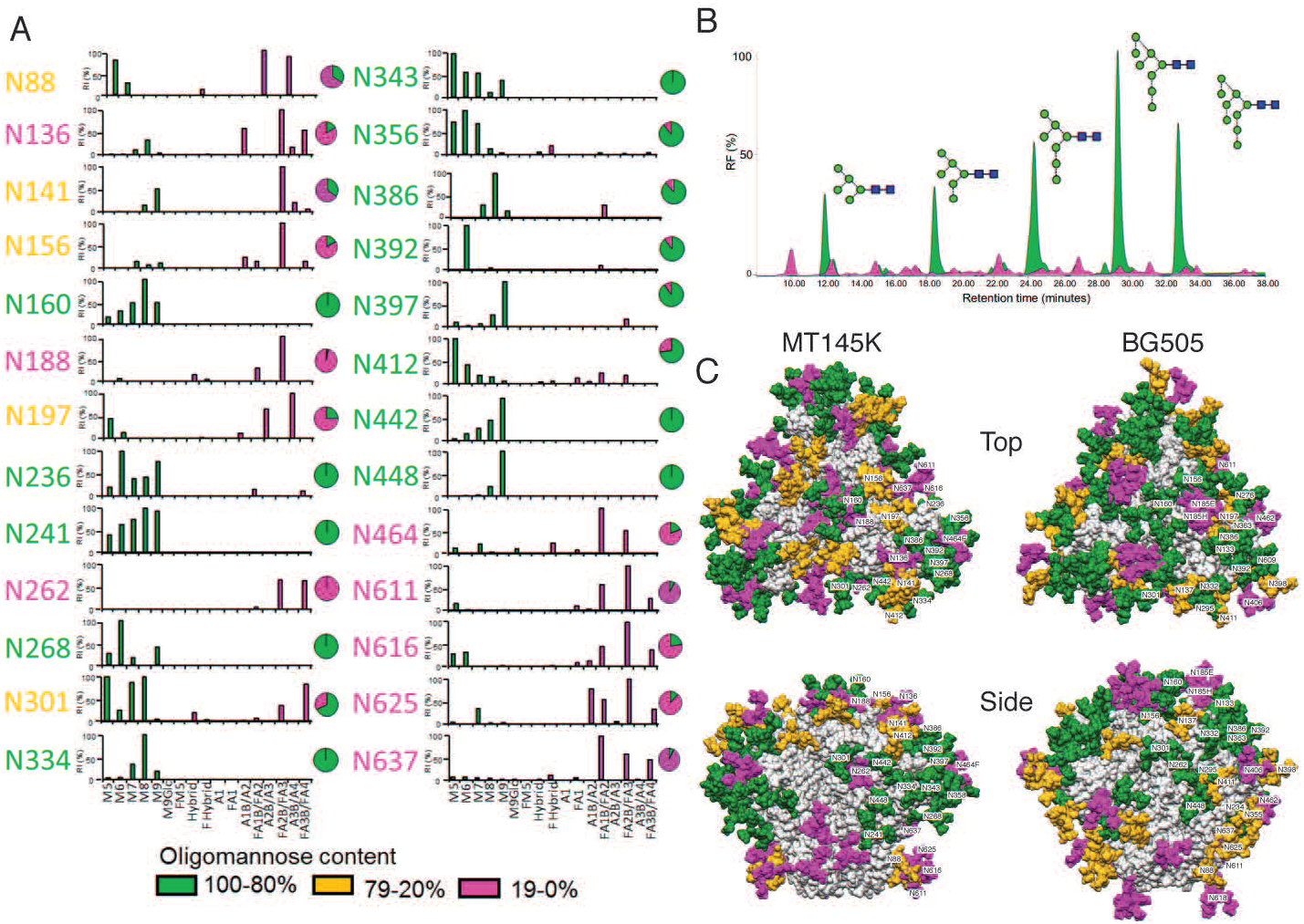
Site-specific glycoform composition of MT145K trimer. A. Site-specific glycoform quantification of the MT145K SOSIP soluble trimer. MT145K trimers from transiently transfected HEK293T cell expressed supernatants were affinity purified by the quaternary trimer-specific antibody, PGT145. The purified MT145K trimers were treated separately with three proteases: trypsin, chymotrypsin and elastase and the digests were enriched for glycopeptides and analysed by LC-ESI MS. The individual glycan composition of the N-linked glycan sites (n=26) is represented by bar graphs that indicate the relative abundance of each glycoform species and are derived from the mean of two analytical replicates. The pie charts summarize the proportion of glycoforms for each site and this information is color coded; oligomannose-type (green) and complex/hybrid glycans (pink). **B.** HILIC-UPLC profiles of the total N-linked glycans released from MT145K trimers. The proportions of oligomannose plus hybrid glycan contents and complex-type glycans are represented in green and pink colors, respectively. **C.** Modeled glycan shields for MT145K and BG505 SOSIP trimers. Man_9_GlcNAc_2_ oligomannose-type glycans were docked and rigid-body-fitted at each of the corresponding Env glycan positions using the MT145K structure (determined in this study (PDB: submit)) and the unliganded BG505 SOSIP.664 trimer structure ((Kwon et al., 2015) PDB: 4ZMJ). Top and side views of the trimers are shown and the individual glycan sites are labelled and color-coded based on the content of oligomannose; green (100-80%), orange (79-20%) and pink (19-0%).

In the MT145K trimer, glycans at N156 and N262 were predominantly complex-type, whereas the corresponding glycans are oligomannose-type in HIV Env (Behrens et al., 2016). These differences may arise due to the proximity of neighbouring glycans. For instance, the HIV Env glycans at positions N295 and N332, adjacent to the N262 glycan, are absent on MT145K Env, which may lead to increased processing of N262 (Figure 3A, C). The remarkable conservation in the overall architecture of the SIV and HIV Env glycan shield, despite sharing only ~62% of the amino-acid sequence identity, suggests that the glycan shield has an indispensable role in immune evasion and potentially maintaining functional integrity of the trimer spike. Indeed, the glycan shield is integral to all lentiviral envelopes and appears to have evolved somewhat specifically to mammalian host (Figure S7). Over the course of lentiviral evolution, the Env glycan density shows an overall gradual progression, and likely peaked in retroviruses infecting non-human primates and plateaued in HIV Envs (Figure S7) (Zhang et al., 2004). Therefore, the high-density Env glycan shield on HIV must have been established well before chimpanzee SIV crossed into humans. Nevertheless, several glycan positions on HIV-1 Env appear to have subtly shifted after the species cross-over that presumably resulted as an adaptation to the human immune system (Figure S8).

### MT145K binds V2 apex bnAbs almost exclusively

To define the overall antigenicity of the MT145K trimer, we first assessed neutralization sensitivity of MT145K virus (MT145-Q171K) to a broad panel of HIV-1 Env-specific neutralizing and non-neutralizing (nnAbs) mAbs and compared these profiles to the clade A BG505 HIV virus (Figure 4A, S9) (Sanders et al., 2013; Voss et al., 2017). Remarkably, the V2 apex bnAbs, but essentially no other bnAbs or nnAbs (except 35O22 gp120-41 interface mAb), exhibited potent neutralizing activities against MT145K virus (Figure 4A, S9). As previously observed, the BG505 isolate was sensitive to neutralization by all of the bnAbs in the panel, but none of the nnAbs (Figure 4A Figure S9).

**Figure 4.**
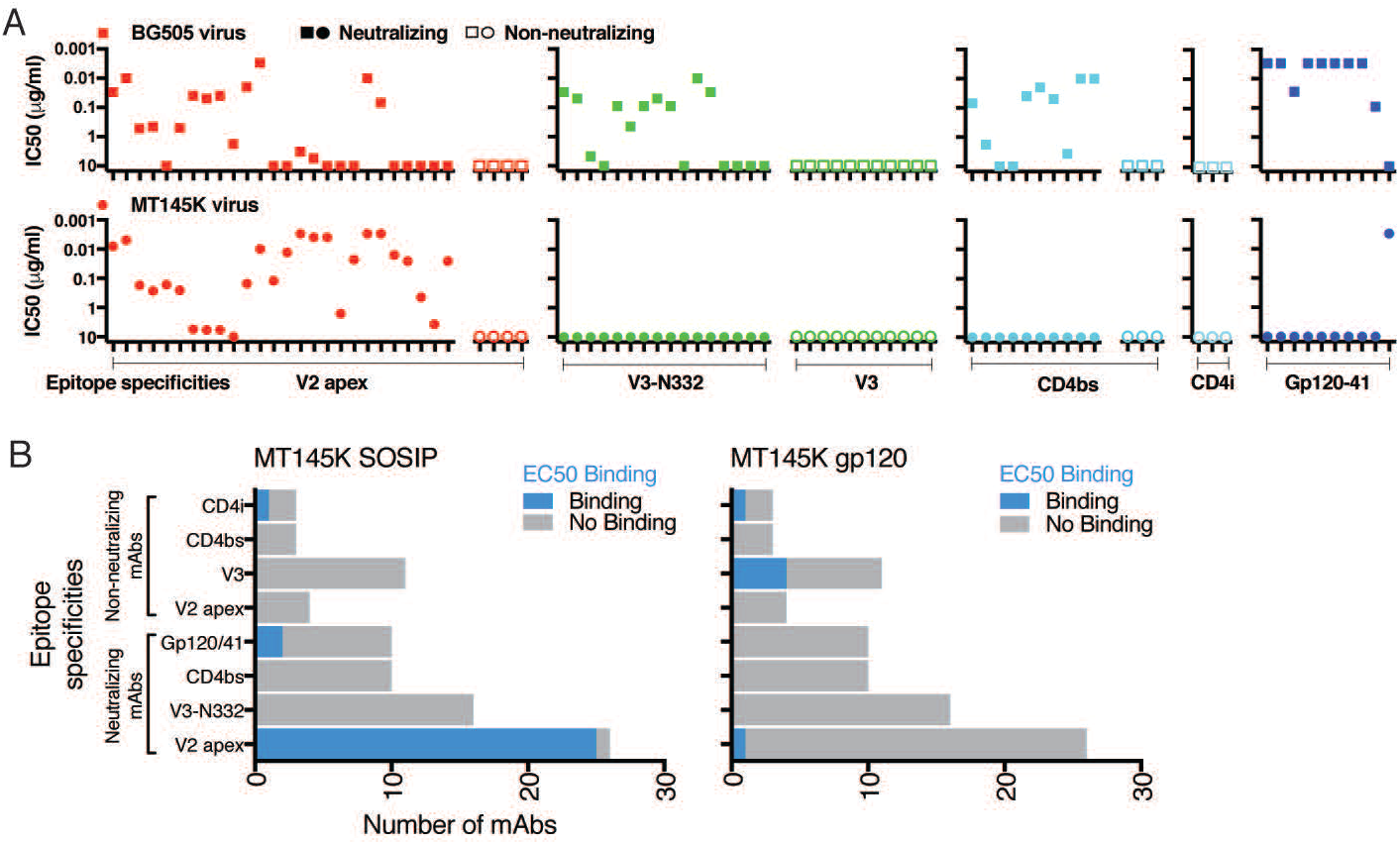
Antigenic profile of the MT145K trimer. **A.** HIV-1 Env-specific mAbs were used to characterize the antigenicity of the MT145K Env trimer. MAbs targeting neutralizing and non-neutralizing epitope specificities, including V2-apex, N332-V3, linear V3, CD4bs, CD4i and gp120-41 interface were tested with MT145K and BG505 Env-encoding pseudoviruses in a TZM-bl cell-based reporter assay. The reciprocal IC_50_ neutralization titers for each virus are indicated as dot plots; plots for individual epitope specificities are depicted separately. The neutralization sensitivity comparison of BG505 and MT145K viruses against the mAb panel shows a selectively potent neutralization of MT145K by V2 apex bnAbs but no other bnAbs, except a single gp120-gp41 interface bnAb, 35022. BG505 virus was neutralized by bnAbs targeting diverse Env sites. **B.** The above mAb panel was further tested with PGT145 Ab-purified MT145K trimer and GNL-purified MT145K gp120 monomer by ELISA. The binding, represented as EC_50_ binding titers, shows selective binding of MT145K by V2 apex bnAbs. Two of the gp120-gp41 interface bnAbs and a CD4i mAb also showed significant binding to MT145K trimer. Four of the non-neutralizing mAbs specific to a linear V3 epitope exhibited binding to MT145K gp120, but not to the trimer.

Next, we evaluated binding of MT145K trimer and monomeric gp120 to a panel of mAbs by ELISA. Consistent with the neutralization results above, bnAbs to the V2 apex site showed robust binding to the MT145K trimer (Figures 4B, S9), but other bnAbs and nnAbs did not bind, except for a few mAbs that displayed very weak binding (Figures 4B, S9). PG9, 17b and some of the linear V3-loop directed mAbs (2557, 3074, 3904 and 14e) (Figures 4B, S9) that bound to the MT145K gp120 monomer. The results suggest that the sequence-dependent epitopes for some of the non-neutralizing V3-loop mAbs are present on monomeric MT145K gp120, but are obscured on the MT145K trimer, as indicated by the MT145K structure. Virus neutralization and trimer binding by mAbs is strongly correlated (p = 0.003), consistent with the notion that the MT145K soluble trimer adopts a native-like trimeric Env configuration and displays antigenic properties optimal for a vaccine immunogen.

### HIV bnAb epitopes on SIV Env

To gain insight into the differences in the HIV-1 Env bnAb epitopes on MT145K SIV Env that may potentially explain the reactivity of V2 apex bnAbs and non-reactivity of HIV bnAbs targeting other Env epitopes, we took advantage of the previously determined structures of human HIV bnAbs in complex with various HIV Env forms and compared the corresponding epitope regions with the MT145K Env (Garces et al., 2014; Lee et al., 2017; Lee et al., 2016; Ozorowski et al., 2017; Pejchal et al., 2011; Wu et al., 2010). A lysine-rich patch in strand C of the V2 loop (^166^RDKKQK^171^ on BG505 Env) and two nearby glycans N160 and N156 form the core epitope for V2 apex bnAbs on HIV Envs (Figure 5A, S10) (Gorman et al., 2016; Julien et al., 2013b; Lee et al., 2017; McLellan et al., 2011; Pancera et al., 2013). Both of these features are conserved on the MT145K trimer, thus enabling the human V2 apex bnAbs to be highly effective against the SIV Envs (Figures 5A, S8, S10) (Barbian et al., 2015).

**Figure 5.**
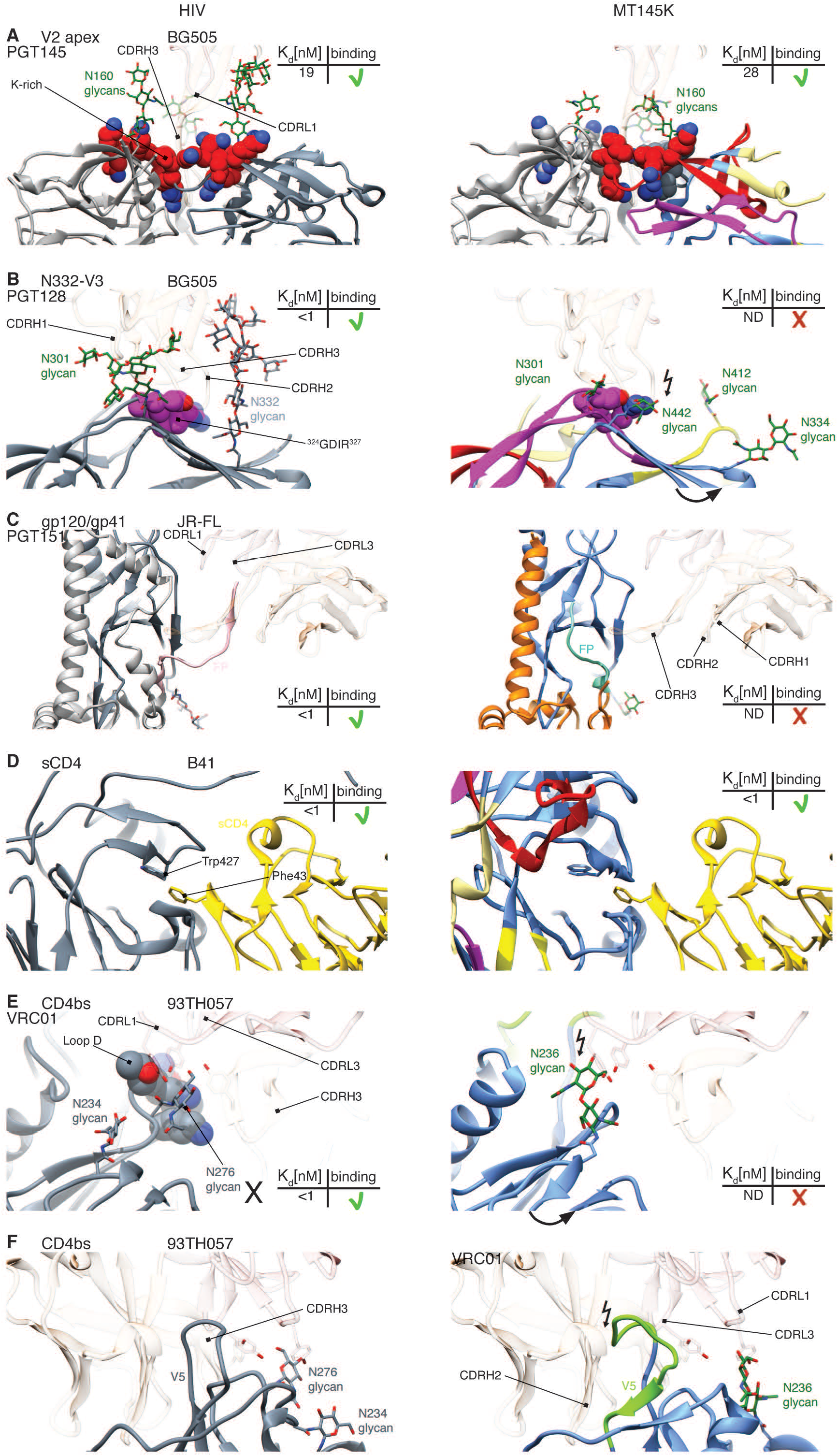
A close-up view of regions on the MT145K trimer that correspond to those recognized by HIV bnAbs on HIV trimers. **A.** V2 apex bnAb binding region: cryo-EM model of PGT145 bnAb (HC: transparent sandy brown; LC: transparent orchid) in complex with BG505 SOSIP trimer depicting V1V2 loops in ribbon representation ((Lee et al., 2017) PDB: 5V8L). The strand C K-rich region (^166^RDKKQK^171^; red spheres) and the glycan N160 (forest green sticks) that form the epitope for PGT145 bnAb are indicated. The elements in the core epitope interact with the CDRL1 loop and the long CDRH3 loop that penetrates through glycans to reach the positively charged surface underneath. Both glycan N160 and the positively charged protein residues are conserved between BG505 HIV-1 and MT145K SIV Env trimers. **B.** V3-glycan bnAb binding region: cryo-EM model of PGT128 bnAb (HC: transparent sandy brown; LC: transparent orchid) in complex with the BG505 SOSIP trimer ((Lee et al., 2015) PDB: 5ACO). The V3 loop protein backbone residues (^324^GDIR^327^; depicted in purple spheres) and the glycans N301 and N332 form the bnAb epitope and are shown to interact with the antibody CDR loops. The MT145K trimer has a glycan at N334 rather than N332 and the glycan points away from the expected location of the PGT128 Ab paratope. In addition, MT145K Env has glycans at two positions N412, (positioned differently on HIV Env) and N442 (absent on HIV Envs) and particularly the latter will clash with PGT128 CDRH2 and prevent it from interacting with the protein part of the epitope. **C.** The gp120-gp41 interface bnAb binding region: cryo-EM model of PGT151 bnAb bound to a membrane-extracted clade B JRFL Env trimer. The structure depicts PGT151 bnAb CDRs interacting with gp120 and the gp41 interface regions ((Lee et al., 2016); PDB: 5FUU). PGT151 CDRH3 interacts with the epitope formed by the protein backbone (in both gp120 and gp41) including the fusion peptide (depicted in pink) and the gp120 (N88, N448) and gp41 (N611 and N637) glycans (not shown). PGT151 Ab CDR loops interact with the FP region on the BG505 trimer. The MT145K trimer FP region (cyan) remains hidden inside the trimer. **D.** Cryo-EM model of 2-domain human sCD4 with B41 SOSIP trimer ((Ozorowski et al., 2017); PDB: 5VN3). The structure shows how the Phe43 residue on sCD4 stacks into the Env cavity lining Trp427. This Trp427 cavity is conserved between HIV-1 and MT145K Envs to accommodate CD4 binding. **E-F.** CD4bs bnAb binding region: crystal structure of VRC01 bnAb in complex with 93TH057 gp120 ((Zhou et al., 2010) PDB: 3NGB). The structure depicts VRC01 CDRH3, CDRL3 and CDRL1 loops interacting with the protein residues in loop D (HXB2: 278-282) and the glycan at N276. The MT145K trimer lacks the N276 glycan and bears glycan N236 (unique to SIV Env) in place of N234 that would clash with the VRC01 CDRL1 loop. Additionally, the MT145K Env trimer has a longer gp120-V5 loop due to a 6-amino acid insertion at HIV HXB2-456 residue that would shift the loop such that it clashes with the CDRH2 the VRC01 Ab.

Binding of one of the N332-V3 epitope specific bnAbs, PGT128, predominantly relies on the N332 glycan and a neighboring peptide motif ^324^GDIR^327^ at the base of the V3 loop (Figures 5B, S10) (Garces et al., 2014; Pejchal et al., 2011; Sok et al., 2016b). The lack of binding to the MT145K trimer by PGT128 and other bnAbs in this class can be explained by the absence of the N332 glycan on this Env. In contrast, 3 of the 4 core protein epitope residues ^324^G-^325^D-^327^R are conserved on MT145K trimer and, in fact, on other chimpanzee SIV Envs (Figures 5B, S8, S10). For the PGT128 class bnAbs, the interaction with glycan N332 can be substituted by the N295 glycan observed in some HIV isolates, but not by glycan N334 that is present on the MT145K trimer (Sok et al., 2014a). In fact, the MT145K N334 glycan points in a different direction away from the N332-V3 epitope site making it impossible to facilitate bnAb binding to this epitope. Strikingly, the majority of known SIVcpz Env sequences possess an N334 glycan in place of the more common N332 glycan on the HIV Env, which appears to be a significant glycan shift upon species cross-over as the virus established itself in humans (Figure S8). In addition, the glycan at N412 in the gp120-V4 region of MT145K Env may obstructively interfere with bnAb binding, and, particularly, the glycan at N442, unique to the MT145K Env trimer and several other SIVcpz Envs, would clash with CDRH2 of PGT128 and other bnAbs in this class and may prevent them from accessing the epitope (Figure 5B, S8).

PGT151 represents another glycan-targeting bnAb class (Blattner et al., 2014; Falkowska et al., 2014; Lee et al., 2016) that recognizes several glycans on gp120 (N88, N448) and gp41 (N611 and N637) as well as the fusion peptide. All glycans and fusion peptide residues that contribute to the PGT151 epitope are conserved between HIV and SIVcpz Envs (Figure S8). Therefore, the lack of PGT151 binding to MT145K is most likely attributable to inaccessibility of the FP on MT145K (Figure 5C).

The CD4bs is conserved between HIV and SIV to the extent that there is cross-species reactivity with sCD4. Human CD4-IgG2 immunoadhesin binds well to the MT145K trimer, indicating a strong cross-species conservation of the Env CD4bs. Phe43 in domain-1 of human sCD4 would fit well inside the Trp427 Env cavity on the MT145K trimer reminiscent of its interaction with the HIV Env BG505 trimer (Figure 5D) (Ozorowski et al., 2017). However, the MT145K trimer is nonreactive with CD4bs bnAbs. VRC01, one of the bnAbs in this class, binds to the HIV Env CD4bs bnAb epitope formed by discontinuous protein backbone elements including loop D of the gp120-C2 region and bordered by a glycan at N276 (Figure 5E-F, S10) (Wu et al., 2010). MT145K lacks the N276 glycan and the proximal N234 glycan, present in most HIV-1 Envs, but instead has a glycan at position 236. Differences in the loop D sequence (Figure S8) and the glycan at N236, which would clash with VRC01 CDRL1 and CDRL3 loops (Figure 5F) on the MT145K trimer likely impose the biggest impediment to VRC01 binding. Further, the MT145K gp120-V5 loop has a 6-amino acid insertion at HXB2 position 456 compared to HIV-1 Envs that would clash with the VRC01 LC (Figure 5F, S8).

Overall, the non-reactivity of HIV Env bnAbs with the MT145K trimer can be largely ascribed to subtle glycan shifts that have occurred in HIV-1 from chimpanzee SIV Env as the virus established itself in humans.

### The engineered MT145K but not the MT145-WT trimer activates V2 apex UCA-expressing B cell precursors *in vivo*

To determine whether the engineered chimpanzee MT145K trimer could efficiently activate HIV V2 apex Ab germline-encoding precursor B cells *in vivo* and how it compares with the MT145-WT trimer, we conducted immunization experiments in the CH01 unmutated common ancestor (UCA) “HC only” knock-in (KI) mouse model. This KI-mouse model expresses the pre-rearranged heavy chain (V_H_DDJ_H_) of the CH01 V2 apex bnAb UCA paired with WT mouse light chains. We immunized two groups of 5 CH01 UCA “HC only” KI mice, each with two repeated doses (at week-0 and week −4) of MT145-WT or MT145K trimer (Figure 6A). To track the development of Ab responses, we performed ELISA assays of the pre-bleed, 2-week (day 14) post prime (Bleed #1) and 2-week post boost-1 (day 42) (Bleed #2) serum samples with MT145K SOSIP trimer protein and its N160 glycan knock-out variant (MT145K N160K) (Figure 6B). The pre-bleed serum samples in both immunization groups exhibited weak binding activity with the MT145K trimer that was dependent on the N160 glycan, consistent with the presence of CH01 UCA Abs that do show some binding to MT145K trimer as described above (Figure 6B).

**Figure 6.**
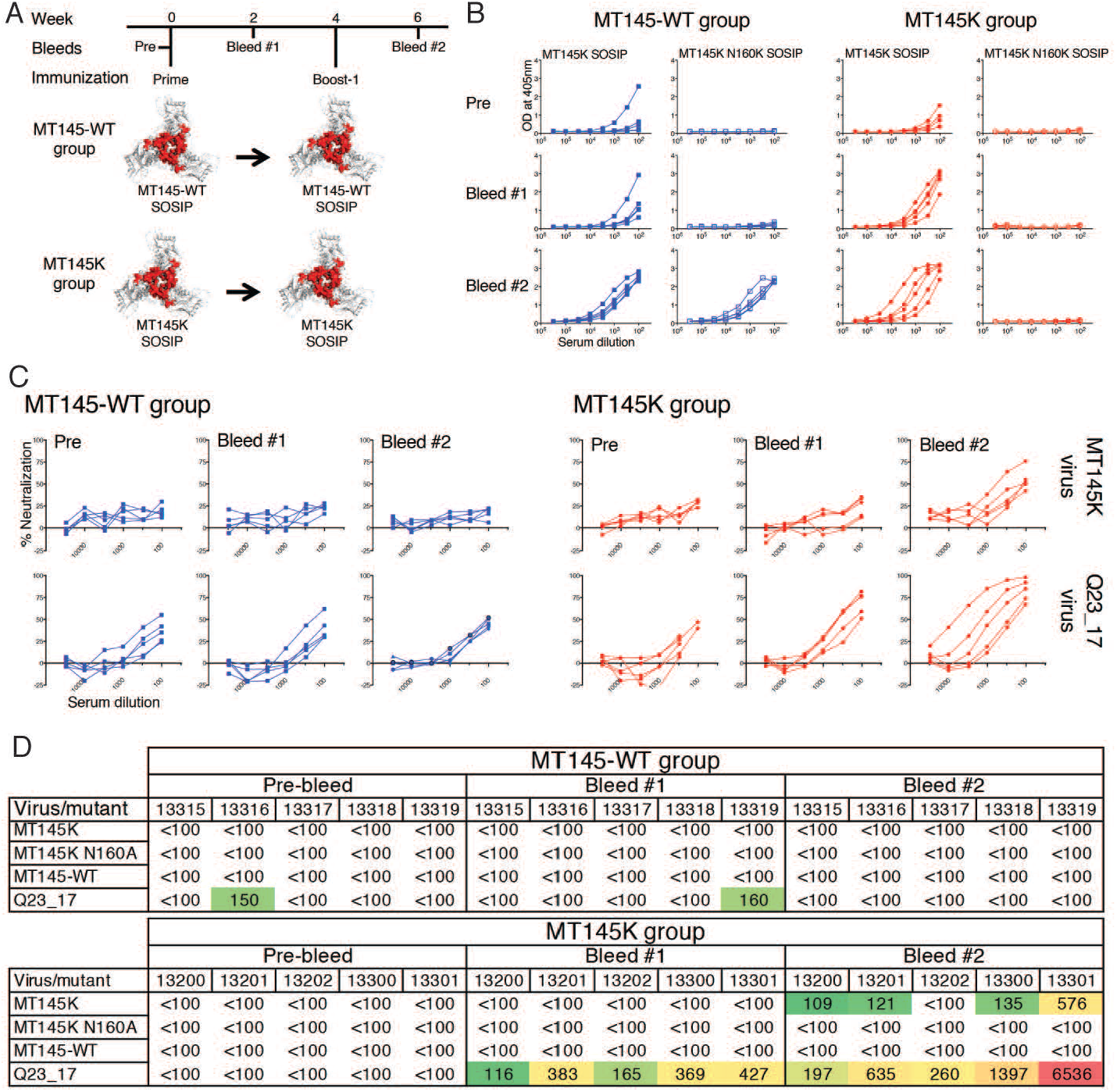
Immunogenicity of MT145-WT compared to MT145K trimers in CH01 UCA HC-only knock-in mice. **A.** Schematic showing immunization schedule of CH01 UCA “HC-only” KI mice with MT145-WT and engineered MT145K trimers. The CH01 UCA “HC-only” KI mice were immunized twice with 25μg of the soluble trimer with GLA-SA as adjuvant. Time points for immunization and bleeds are indicated. **B.** ELISA binding of the MT145-WT and MT145K group trimer-immunized CH01 UCA “HC-only” KI mice serum samples (pre-bleed (Pre), two-weeks post prime (Bleed #1) and two-weeks post boost-1 (Bleed #2)) with soluble MT145K SOSIP and its glycan knock-out variant (MT145K N160K) trimers. **C.** Neutralization titrations of the MT145-WT and MT145K group trimer immunized CH01 UCA “HC-only” KI mice sera (pre-bleed (Pre), post prime (Bleed #1) and post boost-1 (Bleed #2)) with MT145K virus and a CH01-sensitive virus (Q23_17). 3-fold diluted sera were tested against the viruses in a TZM-bl reporter cell assay. **D.** ID_50_ neutralization titers of the MT145-WT and MT145K group trimer-immunized CH01 UCA “HC-only” KI mice sera (pre-and post-immunization bleed time points). Neutralization was assessed against the priming immunogen-matched autologous viruses in each group (MT145-WT group: MT145-WT virus, and MT145K group: MT145K virus), the N160 glycan knock-out variant of MT145K virus (MT145K N160A) and a highly CH01-sensitive virus, Q23_17. The numerical values shown in the table represent the ID_50_ neutralization titers of the immune serum samples and were calculated by non-linear regression method from the percent neutralizations of serum titrations with virus.

The immunogen-specific titers of the serum Ab responses post prime immunizations (Bleed #1 samples) marginally increased in the MT145K group but remained largely unchanged in the MT145-WT trimer immunized group. The serum Ab titers post boost-1 immunization (Bleed #2) increased in both the groups and were orders of magnitude higher as compared to the pre-bleed or the post prime Ab binding responses (Figure 6B). At this immunization step, the serum Ab responses in the MT145K trimer immunized group were solely dependent on the N160 glycan while the MT145-WT trimer immunization group responses targeted the MT145 Env that were mostly independent of the N160 glycan, which forms part of the core V2 apex bnAb epitope (Figure 6B). Therefore, we conclude that the engineered MT145K trimer but not the MT145-WT, efficiently triggers the epitope specific V2-apex bnAb UCA encoding B cell precursors *in vivo*. Remarkably, immunizations with Q171K substituted engineered MT145K trimer also appeared to eliminate the non-V2 apex bnAb site Env specific off-target B cell responses that were elicited in the MT145-WT trimer immunization group (Figure 6B). The results demonstrate that the activation of the HIV Env bnAb-encoding unmutated B cell precursor by immunogens that display binding to their UCA Ab versions is critical for eliciting epitope-specific Ab responses and the findings are consistent with studies that specifically use germline-targeting immunogen molecules to kick-off the bnAb precursor encoding B cell responses *in vivo* (Dosenovic et al., 2015; Escolano et al., 2016; Jardine et al., 2015; McGuire et al., 2013; Sok et al., 2016a; Steichen et al., 2016b; Tian et al., 2016).

Next, we evaluated immune sera for neutralization of autologous and heterologous viruses. Reproducible MT145K autologous virus-specific neutralizing Ab responses were induced in the MT145K immunization group but not in the MT145-WT immunization group (Figure 6C-D). As for the ELISA binding responses, the nAb titers in the MT145K trimer immunized group increased at 2 weeks post prime, as indicated by nAb titers against a highly CH01 sensitive HIV Env-encoding virus (Q23_17), and further significantly increased after the boost-1 immunization (Figure 6C). At this point, all animals in the MT145K group developed autologous virus specific nAb responses (Figure 6C). The nAb responses in MT145K trimer-immunized animals mapped to the glycan N160 and strand C K171 residue, both of which form part of the core epitope for V2-apex bnAbs, suggesting that the MT145K trimer successfully primed V2 apex UCA B cells in an epitope-specific manner *in vivo.*

Overall, we conclude that the engineered MT145K but not the MT145-WT trimer activated the V2-apex specific bnAb precursor B cells in a UCA-expressing mouse model and further drove maturation along favorable B cell pathways.

### Combining chimpanzee SIV MT145K trimer with HIV Env trimer immunizations in the CH01 UCA model

Evaluation of the utility of the MT145K trimer in a sequential HIV immunization regime will be best carried out in humans. Nevertheless, we were interested to investigate the effects of combining the chimpanzee SIV MT145K and HIV Env trimers in a prime-boost immunization in the CH01 UCA “HC only” KI mice. We immunized 4 groups of CH01 UCA KI mice with combinations of chimpanzee SIV MT145K SOSIP and HIV CRF250 SOSIP, previously shown to bind CH01 iGL Ab (Andrabi et al., 2015; Gorman et al., 2016), and finally boosted with an HIV Env derived 3-trimer cocktail (C108, WITO and ZM197-ZM233V1V2 SOSIPs) (Figure 7A). After priming, MT145K trimer-primed animals showed autologous ID50 nAb response in only 1 out of 10 animals while CRF250 priming produced autologous nAb titers in 6 out of the 10 animals (Figure 7B, Table S2). This is the result of the increased sensitivity of the CRF250 virus compared to the MT145K virus to neutralization by CH01 bnAb since both sets of primed animals neutralized the CH01 bnAb-sensitive Q23_17 virus (Figure 7B). The priming also led to development of sporadic cross-neutralizing responses against a few HIV heterologous viruses sensitive to CH01-class bnAbs (Bonsignori et al., 2011), including against the CRF250 virus in MT145K trimer primed animal groups (Figure 7B, Table S2).

**Figure 7.**
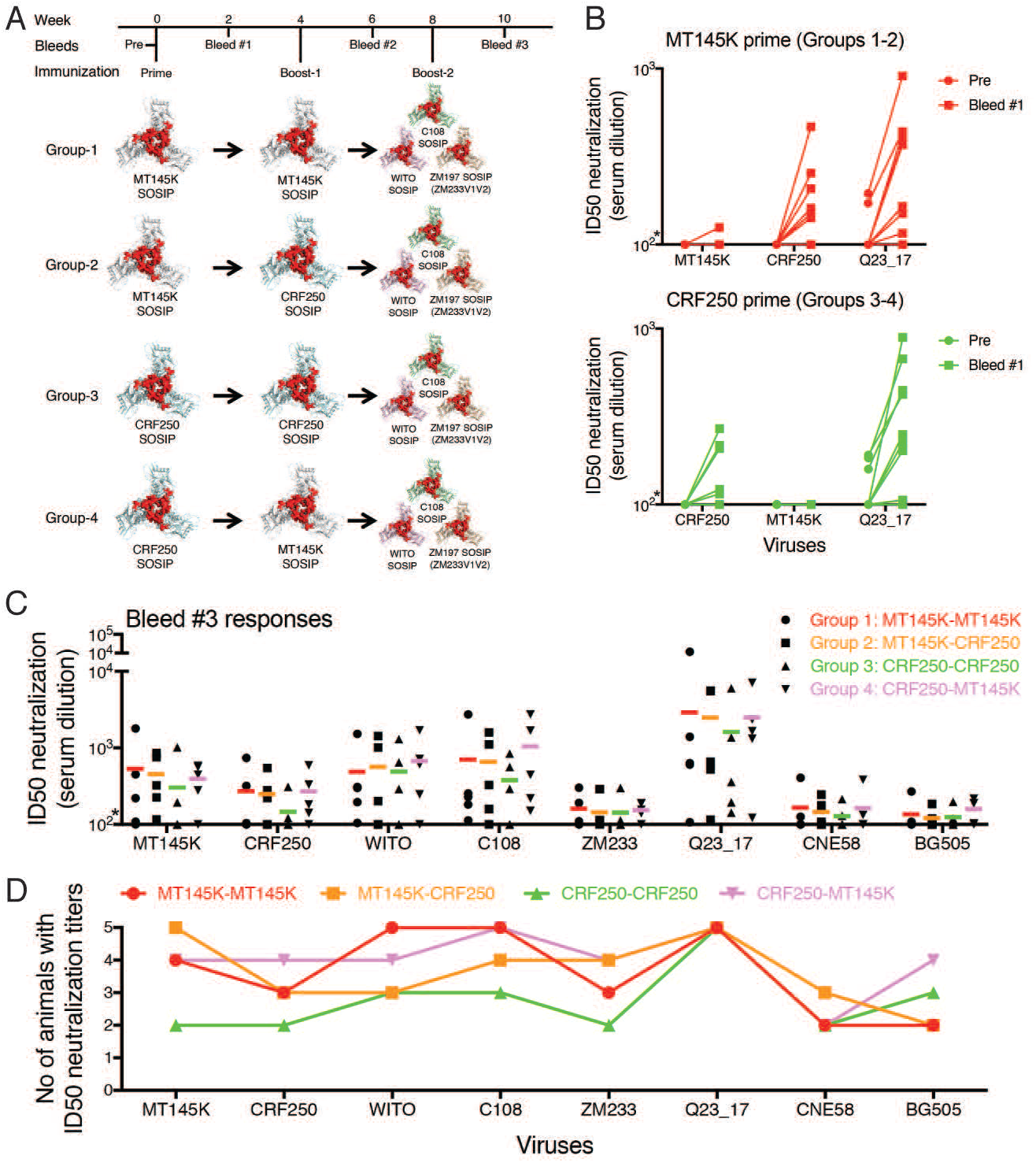
Immunizations combining chimpanzee SIV MT145K trimer with HIV trimers in CH01 UCA HC-only knock-in mice. **A.** Schematic showing immunization schedule of CH01 UCA “HC-only” KI mice with combinations of chimpanzee SIV MT145K trimer and HIV Env derived trimers. Four groups of 5 animals each were immunized with two doses (prime: week-0 and boost-1: week-4) of the trimers as follows; Group-1 (MT145K twice), Group-2 (MT145K followed by CRF250), Group-3 (CRF250 twice) and Group-4 (CRF250 followed by MT145K). Each group was further boosted (boost-2 at week-8) with an HIV Env derived 3-trimer cocktail (C108, WITO and ZM197-ZM233V1V2). The V1V2 loops on trimer cartoons are depicted in red to highlight that the region is shared between HIV and SIV Env trimers. The CH01 UCA “HC-only” KI mice were immunized with 25μg of the soluble trimer (MT145K or CRF250 or HIV trimer cocktail (25μg total)) with GLA-SE as adjuvant. Time points for the immunizations and the bleeds are indicated. **B.** Comparison of CH01 UCA “HC-only” KI mice B cell priming by chimpanzee SIV MT145 trimer and HIV Env derived CRF250 trimer. ID50 neutralization titers of the pre-bleed (Pre) and post-prime (Bleed #1) sera from CH01 UCA “HC-only” KI mice immunized in groups 1 and 2 with MT145K trimer and groups 3 and 4 with CRF250 trimer against MT145K, CFR250 and a highly CH01 sensitive virus Q23_17 are shown. Each dot in the plot represents virus ID50 values for individual animals “*”indicates that 50% neutralization was not reached at a 1:100 serum dilution. **C.** Dot plot showing comparison of ID50 neutralization titers of post boost-2 (Bleed #3) sera from the four different groups of trimer immunized CH01 UCA “HC-only” KI mice against the immunization prime and boost-matched and CH01 sensitive pseudoviruses. Each dot represents an individual ID50 neutralization titer grouped by virus (shown on the x-axis) and the mean ID50 values for each immunization group against each virus (indicated by colored horizontal lines). **D.** The induction of neutralization breadth in CH01 UCA “HC-only” KI mice by the full immunization schedule in Fig 7A. The reproducibility of neutralization breadth is plotted as number of animals in each immunization group (groups indicated by color) that reach 1/ID50>100 against each virus. The Bleed #3 serum ID50 neutralization responses for the groups that received chimpanzee SIV MT145K alone or in combination with HIV CRF250 trimer show a trend for better reproducibility in eliciting neutralization breadth than the group that had HIV Env immunizations. The differences only achieve statistical significance (p=0.045) when comparing reproducibility of neutralization breadth between groups 3 and 4 using nonparametric Mann-Whitney test.

Homologous (MT145K and CRF250 primed animals boosted with MT145K and CRF250, respectively) and heterologous (MT145K and CRF250 primed animals boosted with CRF250 and MT145K, respectively) boosting immunizations produced stronger nAb responses in all the animals suggesting an immune recall response. There was a general improvement in the neutralization breadth against heterologous viruses (Table S2), suggesting that the boosting immunizations resulted in development of B cell responses along favorable maturation pathways. The responses mapped entirely to N160 glycan and the strand C residues that form part of V2 apex bnAb core epitope (Andrabi et al., 2015) (Table S2). Of note, there was no significant difference between homologous and heterologous boosting in this model.

Following further boosting with an HIV trimer cocktail, a number of animals in all the groups developed some neutralization breadth (Fig 7C). There was a trend for greater breadth in the animals in which SIV and HIV trimer immunizations were combined but this only achieved significance (p=0.045) when comparing the CRF250-MT145K Env and the CRF250-CRF250 Env prime/boost regimes. We were confounded that greater differences between SIV/HIV and HIV/HIV immunizations were not observed as might be anticipated by an expected reduction in off-target responses in the former case. However, the frequency of B cell precursors in the mice is very high and this factor may have allowed the HIV/HIV regime to be more effective than it would be when precursor frequencies are much lower by enhancing the likelihood of on-target relative to off-target responses.

Overall, the analysis of the immune responses revealed that, due to the extraordinary conservation of the V2 apex bnAb epitope region between HIV and chimpanzee SIV, the MT145K trimer successfully primed human V2 apex bnAb UCA-encoding mouse B cells and induced a V2-focused cross-neutralizing HIV Env specific response that could be further boosted by HIV Env derived trimers

## Discussion

Vaccination has taken advantage of related viruses from a different species, beginning with the use of cowpox as a smallpox vaccine (Riedel, 2005). HIV is too variable and has too many evasion mechanisms for such an approach applied directly to work effectively. Nevertheless, there are HIV related viruses that have the potential to be exploited in some form in vaccine design. Indeed, the HIV pandemic is believed to have arisen because of a cross-species virus transmission from chimpanzees to humans in the period from 1910-1930 (Korber et al., 2000; Sharp and Hahn, 2011; Worobey et al., 2008). The HIV and chimpanzee SIV Envs, the target of potentially protective neutralizing antibodies, display about 60% sequence conservation at the amino acid level. Importantly, HIV V2-apex bnAbs have been shown to neutralize certain chimpanzee SIV isolates, including the SIVcpz*Ptt* isolate MT145, suggesting cross-species conservation of this epitope. Accordingly, we generated a chimpanzee SIV Env trimer (MT145 SOSIP) and showed that it bound HIV V2-apex bnAbs. We then engineered it to bind to germline-reverted V2 apex bnAbs (MT145K SOSIP) so that it might be useful in activating V2-apex precursors.

The cryoEM structure of MT145K SOSIP trimer revealed that the Env trimers of HIV and chimpanzee SIV have very similar overall architectures. The glycan shield of chimpanzee SIV forms a similarly dense protective layer to antibody recognition of the protein surface as observed in HIV. However, subtle movements in the locations of the glycans contribute to the inability of the great majority of HIV-1 bnAbs to recognize the chimpanzee SIV Env trimer. As noted above, bnAbs to the V2 apex region of the trimer are the exception. We have hypothesized previously (Lee et al., 2017) that the conservation of this region amongst HIV isolates is to facilitate trimer disassembly during viral entry. It is interesting that the overall V2 apex structure is conserved across the chimpanzee-human species barrier indicating its critical importance for Env function.

In order to evaluate, MT145K trimer as an immunogen able to activate V2-apex bnAb precursor B cells, we took advantage of the availability of V2-apex bnAb UCA H chain only knock-in mice. We compared MT145K and MT145 trimers as immunogens. Following two immunizations, MT145K trimers reproducibly elicited Abs able to neutralize the autologous virus and a few V2-apex Ab sensitive viruses whereas MT145 trimers failed to induce such nAbs. The specificities of the nAbs were dependent on the glycan at N160 and a lysine on strand C of the V2. Boosting with a cocktail of HIV Env trimers successfully recalled the V2 apex specific nAb responses and generated some enhanced heterologous neutralization. Therefore, from studies in this mouse model, the MT145K trimer appears a promising immunogen.

In conclusion, chimpanzee SIV Env trimers closely resemble HIV Env trimers with key differences that likely reflect the different immune pressures exerted by the human compared to the chimpanzee antibody repertoire. Nevertheless, the retention of the V2-apex bnAb region and its behavior in a mouse model suggests that the chimpanzee SIV Env can find application in sequential HIV vaccination strategies.

## STAR+METHODS

Detailed methods are provided in the online version of this paper and include the following:

- METHOD DETAILS

- SIV envelope trimer design, its expression and purification
- Antibodies, expression and purification
- Site-directed mutagenesis
- Differential Scanning Calorimetry
- Negative stain electron microscopy and data treatment
- CryoEM sample preparation, data collection, processing and analysis
- Model building and refinement
- Global N-linked glycan analysis
- LC-MS glycopeptide analysis
- Glycan modeling
- Pseudovirus production
- Neutralization assay
- ELISA binding assay
- Bio Layer Interferometry (BLI) binding assay
- Trimer protein immunizations in CH01 UCA HC-only KI-mice
- Data availability

## AUTHOR CONTRIBUITIONS

R.A., J.P., J.A., J.Z., L.V., A.B.W., and D.R.B. designed the experiments. R.A., J.P., J.A., G.S., J.Z., N.D.V., G.G., K.P., C.Y.S., M.P., A.N., and F.G. performed the experiments. H.B.V., I.A.W., M.C., B.H.H., and B.F.H. contributed critical reagents. R.A. J.P., J.A., A.B.W., and D.R.B. analyzed the data and wrote the paper, with inputs from other authors. R.A. and D.R.B. conceived the idea of using SIVcpz*Ptt* Env-derived trimer as an HIV vaccine template.

## ACKNOWLEDGMENTS

This work was supported by the International AIDS Vaccine Initiative (IAVI) through Neutralizing Antibody Consortium SFP1849 (D.R.B., A.B.W., I.A.W.); the National Institute of Allergy and Infectious Diseases (Center for HIV/AIDS Vaccine Immunology and Immunogen Discovery Grant UM1AI100663) (to D.R.B., A.B.W., I.A.W.), the Ragon Institute of MGH, MIT, and Harvard (D.R.B.). This study was made possible by the generous support of the Bill and Melinda Gates Foundation Collaboration for AIDS Vaccine Discovery (CAVD, OPP115782 and OPP1084519) and the American people through USAID. We thank Christina Corbaci and for her help in the preparation of figures.

## METHOD DETAILS

### SIV envelope trimer design, its expression and purification

SOSIP.664 HIV-1 Env trimer modification were incorporated into envelope encoding sequences corresponding to four Chimpanzee (SIVcpz*Ptt*) isolates (GAB1 [GenBank: P17281]; MB897 [GenBank: ABU53023]; EK505 [GenBank: ABD19499]; and MT145 [GenBank: ABD19508]) to express as soluble native trimers as described previously (Sanders et al., 2013). Briefly, the following modifications were incorporated into these Envs for soluble trimer expression: a) the Env leader sequence was replaced by Tissue Plasminogen Activator (TPA) signal sequence for higher protein expression; b) a disulfide bond was introduced between gp120 and gp41 subunits by substituting residues A501-C and T605-C respectively in gp120 and gp41; c) the gp120 REKR cleavage site was replaced by Furin inducible R6 site (RRRRRR) for enhancing cleavage efficiency between gp120 and gp41; and d) an I559P substitution in gp41 to stabilize the soluble trimer protein. In addition, a GS-linker and a His-tag were added to the gp41_ECTO_ C-terminus at HXB2 residue 664 position. The codon-optimized SOSIP.664 gp140 gene constructs were synthesized (Geneart, Life Technologies) and cloned into the phCMV3 vector (Genlantis). Recombinant envelope proteins were expressed in HEK293F cells as described elsewhere (Sanders et al., 2013). Briefly, HIV-1 Env trimers CRF250, WITO, C108, ZM197-ZM233V1V2 and the 4 chimpanzee SIV SOSIP.664 Env-encoding trimer plasmids were cotransfected with a plasmid encoding for Furin (3:1 ratio) into HEK293F cells using PEI-MAX 4000 transfection reagent (*Polysciences, Inc.*). The secreted soluble trimers proteins were purified from cell supernatants after 5 days using agarose-bound Gallanthus Nivalis Lectin (GNL) (Vector Labs) or CNBr-activated Sepharose 4B bead (GE Healthcare) bound PGT145 bnAb antibody affinity columns as described previously (Pugach et al., 2015). The affinity-purified proteins were size exclusion chromatography (SEC)-purified with a Superdex 200 10/300 GL column (GE Healthcare) in PBS/TBS. The purified trimers for the immunization experiments were quality control tested for antigenicity with a range of HIV-1 Env-specific neutralizing and non-neutralizing mAbs.

### Antibodies, expression and purification

HIV-1 envelope specific mAbs to a broad range of epitopes were used, including those that target V2-apex, V3-N332, linear V3, CD4bs, CD4i and gp120-41 Env sites. A dengue antibody (DEN3) was used as control Ab for binding experiments. For PG9 and CH01 V2-apex bnAb inferred germline antibody designs, the heavy and the light chain V-gene of the mature Abs were reverted to their corresponding closest inferred germline gene sequence as determined using the ImMunoGeneTics (IMGT) website (http://imgt.cines.fr/) (Brochet et al., 2008). The reverted variable heavy and light chain nucleotide sequences were synthesized by Geneart (Life Technologies) and cloned into corresponding Igγ1, Igκ, and Igλ expression vectors as previously described (Tiller et al., 2008), using the Gibson cloning method (NEB, USA). The antibodies were expressed and purified using methods described previously (Sok et al., 2014b). Briefly, the heavy and light chain encoding plasmids were reconstituted (1:1 ratio) in Opti-MEM (Life Technologies), and cotransfected HEK293F cells (Invitrogen) using 293fectin (Invitrogen). The suspension cells were cultured for 4-5 days in a shaker incubator at 8% CO2, 37.0°C, and 125 rpm. The antibody containing supernatants were harvested, filtered through a 0.22 mm Steriflip units (EMD Millipore) and passed over a protein A or protein G affinity column (GE Healthcare). The bound antibody was eluted from the columns in 0.1 M citric acid, pH 3.0. Column fractions containing IgG were neutralized (2M Tris-base), pooled, and dialyzed against phosphate-buffered saline (PBS), pH 7.4. IgG purity was determined by sodium dodecyl sulfate-polyacrylamide gel electrophoresis, and the concentration was determined by measuring the relative absorbance at 280 nm.

### Site-directed mutagenesis

The amino-acid point mutations in Env-encoding plasmids were incorporated by using a QuikChange site-directed mutagenesis kit (Agilent Technologies, USA), according to the manufacturer’s instructions. All of the mutations were confirmed by DNA sequence analysis (Eton Bioscience, San Diego, CA).

### Differential Scanning Calorimetry

Thermal denaturation was analyzed with a differential scanning calorimetry (DSC) using a MicroCal VP-Capillary DSC instrument (Malvern), at a scanning rate of 1 K/min under 3.0 atmospheres of pressure. Samples were dialyzed in PBS pH 7.4 overnight and protein concentration was adjusted to 0.5 mg/mL prior to measurement. DSC data were analyzed after buffer correction, normalization, and baseline subtraction using MicroCal VP-Capillary DSC analysis software provided by the manufacturer.

### Negative stain electron microscopy and data treatment

Purified M145K sample was deposited on thin-carbon-coated (Edwards Auto 306 carbon evaporator) a C-flat EM grid (Cu400 mesh, 2μm hole diameter, 2μm hole spacing) (Protochips, Morrisville, NC, USA) and embedded in 2% (w/V) uranyl formate. The carbon-coated grids were Ar/O_2_-plasma-cleaned (Gatan Solarus Model 950 Advanced Plasma System; Gatan Inc., Pleasanton, CA, USA) prior to sample deposition. The uranyl-stained EM sample was then inserted into an FEI Tecnai 12 microscope (Thermo Fisher Scientific, Waltham, MA, USA) equipped with a US4000 CMOS detector (Gatan Inc., Pleasanton, CA, USA). The data was collected at 52,000X nominal magnification resulting in a pixel size of 2.05Å at the object level. Data was binned by a factor of 2 prior to data treatment. Projection image identification in the micrographs was performed with a difference-of-Gaussians implementation (Voss et al., 2009). Projection images subsequently underwent 2D alignment and classification by iterative multi-reference alignment/multivariate statistical analysis (Ogura et al., 2003).

### CryoEM sample preparation, data collection, processing and analysis

Purified MT145K sample was deposited on a C-flat EM grid (Cu400 mesh, 2μm hole diameter, 2μm hole spacing) (Protochips, Morrisville, NC, USA) that had been Ar/O_2_-plasma-cleaned (Gatan Solarus Model 950 Advanced Plasma System; Gatan Inc., Pleasanton, CA, USA) prior to sample deposition. Excess buffer was then blotted away from the grid followed by plunging into and vitrification in liquid ethane cooled by liquid nitrogen using a vitrobot (Thermo Fisher Scientific, Waltham, MA, USA). The resulting cryo-EM specimen was transferred into an FEI Titan Krios microscope (Thermo Fisher Scientific, Waltham, MA, USA) equipped with a Gatan K2 Summit direct electron detector (Gatan Inc., Pleasanton, CA, USA). Dose-fractionated data was collected in electron counting mode at a nominal magnification of 29,000X resulting in a pixel size of 1.02 Å at the object level. Micrograph movie frame exposure time was 200ms and each movie micrograph was recorded over 10s (50 movie frames) corresponding to a total dose of 94e^-^/Å^2^. Movie micrograph frames were subsequently aligned (MotionCor2; (Zheng et al., 2017)), dose-weighted and signal-integrated resulting in 1,281 micrographs for further data processing. CTF models were determined using GCTF (Zhang, 2016). Candidate projection images of MT145K were identified using a difference-of-Gaussians implementation (Voss et al., 2009). The resulting set of candidate projection images subsequently underwent 2D alignment and classification by use of Relion 2.1b1 (Scheres, 2012). ~95,000 projection images corresponding to well-formed class averages of MT145K were selected for further data processing. This data class was iteratively angularly refined and reconstructed using a B41 unliganded Env trimer map rendered at 60 Å resolution as an initial reference (Ozorowski et al., 2017). The data class then underwent 3D classification into six classes with the initial reconstruction rendered at 60 Å resolution as reference. From 3D classification, a subset of 44,301 projection images was selected for final data processing comprising CTF model adjustment at the projection-image level (Zhang, 2016) and angular refinement and reconstruction (Scheres, 2012).

### Model building and refinement

A homology model (Modeller; (Webb and Sali, 2016)) was generated from sequence alignment of MT145K and BG505 and the structure of the latter (PDB: 4TVP). Significant manual rebuilding followed in Coot (Emsley and Cowtan, 2004). A fragment library was then created from the MT145K sequence containing 200 homologous, non-redundant sequences at each MT145K 7-mer position. Library fragment-based, density-guided, real-space rebuilding was then performed (DiMaio et al., 2015) with 319 decoys. The resulting models were evaluated geometrically (MolProbity; (Chen et al., 2010)) and by fit-to-map (EMRinger; (Barad et al., 2015). The overall best model was selected for further iterations of manual rebuilding and multi-decoy, density-guided, real-space, all-atom Rosetta FastRelax refinement. Finally, glycans were manually built in Coot and restricted, density-guided real-space refinement performed in Phenix 1.12 (Adams et al., 2002) followed by model evaluation by MolProbity, EMRinger and Privateer (Agirre et al., 2015).

### Global N-linked glycan analysis

The quantifications and structural characterization of the total glycan pool was achieved by cleaving the N-inked glycans from the surface of the glycoprotein using an in-gel digestion with peptide N-glycosidase F (PNGaseF). The resultant glycans were separated into two aliquots. The first was derivatized with 2-aminobenzoic acid (2-AA) and subjected to HILIC-UPLC analysis using an Acquity UPLC (Waters). To quantify the oligomannose content of the released glycans, the labelled samples were treated with endoglycosidase H (endoH), which selectively cleaves oligomannose glycans. Data analysis and interpretation were performed using Empower software(Waters). The second aliquot of released glycans was subjected to negative ion electrospray ion mobility mass spectrometry using a Synapt G2Si mass spectrometer (Waters). Glycan compositions were determined using collision induced dissociation (CID) fragmentation. Data analysis was performed using Waters Driftscope (version 2.8) software and MassLynxTM (version 4.1). Spectra were interpreted as described previously (Harvey et al., 2009). The glycan compositions were used to generate a sample-specific glycan library that was used to search the glycopeptide data to minimize the number of false-positive assignments in site-specific analysis.

### LC-MS glycopeptide analysis

Site-specific N-glycosylation analysis was performed using proteolytic digestion followed by tandem LC-MS. Prior to digestion, trimers were denatured, reduced and alkylated by incubation for 1h at room temperature (RT) in a 50 mM Tris/HCl, pH 8.0 buffer containing 6 M urea and 5 mM dithiothreitol (DTT), followed by the addition of 20 mM iodacetamide (IAA) for a further 1h at RT in the dark, and then additional DTT (20 mM) for another 1h, to eliminate any residual IAA. The alkylated trimers were buffer-exchanged into 50 mM Tris/HCl, pH 8.0 using Vivaspin columns (GE healthcare) and digested separately with trypsin, elastase and chymotrypsin (Mass Spectrometry Grade, Promega) at a ratio of 1:30 (w/w). Glycopeptides were selected from the protease-digested samples using the ProteoExtract Glycopeptide Enrichment Kit (Merck Millipore) following the manufacturer’s protocol. Enriched glycopeptides were analyzed by LC-ESI MS on an Orbitrap fusion mass spectrometer (Thermo Fisher Scientific), as previously described (Behrens et al., 2016), using higher energy collisional dissociation (HCD) fragmentation. Data analysis and glycopeptide identification were performed using ByonicTM (Version 2.7) and ByologicTM software (Version 2.3; Protein Metrics Inc.), as previously described (Behrens et al., 2016).

### Glycan modeling

Man_9_GlcNAc_2_ oligomannose-type glycans were docked and rigid-body fitted at each of the corresponding Env glycan positions using the MT145K structure presented here or an unliganded BG505 SOSIP.664 structure (PDB: 4ZMJ).

### Pseudovirus production

To produce pseudoviruses, Env-encoding plasmids were cotransfected with an Env-deficient backbone plasmid (pSG3ΔEnv) (1:2 ratio) using X-tremeGENE™ 9 (Sigma-Aldrich) DNA transfection reagent. Briefly, 1X10^6^ cells in 10ml of Dulbecco’s Modified Eagle Medium (DMEM) were seeded in a 100mm x 20mm cell culture dish (Corning) one day prior to transfection. For transfection, 40μl of X-tremeGENE™ 9 was added to 700μl of Opti-MEM I reduced serum medium (Thermo Fisher) in tube 1. The Env-encoding plasmid (5 μg) and pSG3ΔEnv (10 μg) were added to tube 2 in 700μl of Opti-MEM. The tube 1 and tube 2 solutions were mixed together and incubated for 25 min at room temperature. Next, the transfection mixture was added to the media with 293T cells seeded previously and then distributed uniformly. All pseudoviruses were harvested 48-72 h posttransfection, filtered through 0.22 mm Steriflip units (EMD Millipore) and aliquoted for use in neutralization assays.

### Neutralization assay

Neutralization was measured by using single-round replication-defective HIV Env-pseudoviruses and TZM-bl target cells (Montefiori, 2005; Seaman et al., 2010). 25ul of 3-fold serially diluted mAbs or serum samples were pre-incubated at 37°C for 1h with 25ul of tissue culture infective dose-50 (TCID_50_) Env-pseudotyped virus in a half-area 96-well tissue culture plate. TZM-bl target cells (5,000 cells/well) in 50μl of DMEM were added and the plates were allowed to grow in humidified incubator at 37°C and 5% Co_2_. The luciferase activity of the lysed cells was read on instrument (Biotek) after 2-3 days, by adding lysis buffer followed by Brightglow (Promega). The 50% inhibitory concentration (IC_50_) or 50% inhibitory doses (ID_50_) was reported as the antibody concentration or serum dilution required to reduce infection by half.

### ELISA binding assay

ELISA binding experiments were performed as described previously with minor modification (Sanders et al., 2013). ELISA binding with SOSIP.664 trimer proteins with mAbs was carried out by either capturing the trimer proteins onto the anti-His capture antibodies or on the streptavidin coated plates through biotinylated trimers. For trimer biotinylation, the SOSIP.664 proteins were randomly biotinylated using a 2:1 molar ratio of biotin reagent to trimer using the EZ-link-NHS-PEG4-Biotin kit (Thermo Fisher Scientific, 21324). MaxiSorp plates (Thermo Fisher Scientific) were coated overnight at 4C with 2 ug/mL of anti-His Ab (Thermo Fisher Scientific) or 2 ug/mL streptavidin (Thermo Fisher Scientific). Plates were blocked for 1 hr with 3% BSA and washed three times with 0.05% Tween 20-PBS (PBS-T) (pH 7.4). Anti-His or Streptavidin-coated plates were incubated with biotinylated trimers in 1%BSA plus PBST for 1.5 hr and washed three times with PBST. 3-fold serially diluted mAbs or sera were added starting at a maximum concentration of 10 ug/mL (100ug/ml for iGL Abs) (sera at 1:100 dilution) in 1% BSA plus PBST, and incubated at room temperature (RT) for 1.5 hr. Plates were washed three times with PBST. Alkaline-phosphatase-conjugated goat anti-human IgG Fc secondary antibody (Jackson ImmunoResearch Laboratories) was diluted 1:1000 in 1% BSA PBST and added to plates for 1 hr at RT. Plates were washed three times with PBST and incubated with phosphatase substrate (Sigma) for 15 mins and the absorbance at 405 nm recorded. The 50% binding (EC_50_) was recorded as the half of the maximum binding activity and was calculated by linear regression method using Prism 6 Software.

### Bio Layer Interferometry (BLI) binding assay

The binding experiments of Abs to the affinity purified trimers were performed with an Octet K2 system (ForteBio, Pall Life Sciences). Briefly, the mAbs or IgGs (10 ug/mL in PBST) were immobilizing onto hydrated anti-human IgG-Fc biosensors (AHC: ForteBio) for 60 seconds to achieve a binding response of at least 1.0. After Ab capture, the sensor was placed in a PBST wash buffer to remove the unbound Ab to establish a baseline signal. Next, the IgG immobilized sensor was dipped into a solution containing SOSIP.664 trimer protein as analyte and incubated for 120 seconds at 1000 rpm. Following this, the trimer bound to IgG immobilized sensor was removed from the analyte solution and placed into the PBST buffer for 240 seconds at 1000 rpm. The 2 and 4 minute binding intervals respectively denote the association and dissociation binding curves reported in this study. The sensograms were corrected with the blank reference and fit (1:1 binding kinetics model) with the ForteBio Data Analysis version.9 software using the global fitting function. The data are represented as maximum binding response or the association and dissociation curve fits.

### Trimer protein immunizations in CH01 UCA HC-only KI-mice

For the immunization experiments, groups of 5 CH01 UCA HC-only knock-in B cell expressing mice were immunized with 25ug of the individual trimer protein or 25ug total protein of the 3-trimer cocktail (Prime, week-0; Boost-1, week-4 and Boost-2, week-8) along with Glucopyranosyl Lipid Adjuvant in stable emulsion (GLA-SE) as adjuvant. Immunizations were administered intramuscular in the leg of each animal with 25μg of total trimer immunogens. Blood samples were collected at pre-bleed, 2-weeks each, post-prime (Bleed #1), post boost-1 (Bleed #2) and post boost-2 (Bleed #3) immunization time-point for the isolation of sera that were tested for presence of neutralizing antibodies in TZM-bl cell based assay. Serum samples were heat inactivated for potential complement activity at 56°C for 0.5 h. Mice used in this study were approved by Duke University Institutional Animal Care and Use Committee-approved animal protocols.

### Data availability

Cryo-EM reconstructions have been deposited in the Electron Microscopy Data Bank under the accession numbers XXX.

